# Role of the central junction in folding topology of the protein-free human U2-U6 snRNA complex analyzed by time-resolved FRET

**DOI:** 10.1101/825547

**Authors:** Huong Chu, Nancy L. Greenbaum

**Affiliations:** Dept of Chemistry, Hunter College of The City University of New York, New York NY 10065; Ph.D. Program in Chemistry, The Graduate Center of the City University of New York, New York, NY 10016; Ph.D. Program in Biochemistry, The Graduate Center of The City University of New York, New York NY 10016

**Keywords:** spliceosome, U2-U6 snRNA complex, fluorescence, time-resolved FRET, junction

## Abstract

Intron removal during splicing of precursor pre-mRNA requires assembly of spliceosomal small nuclear (sn)RNAs into catalytically competent conformations to promote two transesterification reactions. U2 and U6 snRNA are the only snRNAs directly implicated in pre-mRNA splicing catalysis, but rearrangement and remodeling steps prior to catalysis require numerous proteins. Previous studies have shown that the protein-free U2-U6 snRNA complex adopts two conformations characterized by four and three helices surrounding a central junction in equilibrium. To analyze the role of the central junction in positioning the two helices critical for formation of the active site, we used ensemble time-resolved fluorescence resonance energy transfer to measure distances between fluorophores at selected locations in constructs representing the protein-free human U2-U6 snRNA complex. Data describing four angles in the four-helix conformer suggest the complex adopts a tetrahedral geometry; addition of Mg^2+^ results in shortening of the distances between neighboring helices, indicating compaction of the complex around the junction. In contrast, the three-helix conformer shows a closer approach between the two helices bearing critical elements, but addition of Mg^2+^ widens the distance between these stems. Presence of Mg^2+^ also enhances the steady state fraction of the three-helix conformer found to be active in spliceosomes. Although the central junction assumes a significant role in orienting helices, in neither conformer, with or without Mg^2+^, are the critical helices positioned sufficiently close to favor interaction, implying that a major role of spliceosomal remodeling proteins is to overcome such distances to create and stabilize a catalytically active fold.

## INTRODUCTION

The splicing of precursor messenger (pre-m)RNA is a fundamental biological process in which noncoding RNA segments (introns) are excised and flanking coding regions (exons) ligated to create a contiguous sequence of exons to be translated into protein. This process is catalyzed by the spliceosome, a huge and dynamic ribonucleoprotein complex located in a eukaryotic cell’s nucleus involving five small nuclear (sn)RNAs (U1, U2, U4, U5 and U6) associated with specific proteins to form small nuclear ribonucleoprotein particles (snRNPs and numerous non-snRNP splicing factors) at different stages of assembly. The spliceosome promotes splicing through two transesterification steps (rev. by Will and Lührmann 2011): in the first step, the 2′-OH of a specific adenosine residue of the intron, called branch site, attacks the 5′ of splice site, resulting in release of the first exon and an intron-3′-exon intermediate as a 2′-3′-5′ branched lariat intermediate. In the second step, the newly freed 3′-OH of the 5′ exon attacks the 3′ splice site, ligating the exons and releasing the lariat intron. These reactions were proposed to involve a two-metal ion center (Steitz and Steitz 1993), in which one Mg^2+^ ion activates the 2′OH branch site nucleophile and the other stabilizes the oxyanion leaving group.

The protein-assisted assembly of a spliceosome on a pre-mRNA substrate involves a highly choreographed set of well-defined events in which snRNPs and protein splicing factors, some partially pre-assembled (Maroney et al. 2000; Stevens et al. 2002), are escorted to the assembling spliceosome or are released. Major conformational changes leading to discrete stages of assembly are mediated by eight helicases and numerous non-ATP-dependent remodeling proteins. Individual steps involve recognition of the 5′ splice site and branch site by the U1 and U2 snRNPs, respectively, followed by introduction of remaining U5, U6 and U4 snRNP as a tri-snRNP (reviewed in Will and Lürhmann 2011). A major rearrangement catalyzed by a helicase leads to pairing between U2 and U6 snRNAs and positioning of the 5′ and 3′ splice sites by the U5 snRNP, accompanied by release of U1 and U4 snRNP to achieve an activated state (B^act^). Subsequent helicase-mediated adjustments bring catalytic RNA components into proximity and proteins associated with the Prp19 complex (NTC) lead to local remodeling to activate the branching step (Chan et al. 2003; Makarova et al. 2004), followed by additional rearrangements, the second catalytic step, and finally dissociation of snRNPs. Thus, the ultimate goal of each of these protein-assisted stages is to usher new components to the assembly, catalyze RNA-RNA rearrangements, and to protect, expose, or juxtapose catalytic elements for catalysis at the appropriate time. Detailed images of spliceosomes trapped in defined stages produced by cryo-EM studies have contributed dramatically to our understanding of the landscape, interactions, and rearrangements of spliceosomal components throughout the cycle of assembly and activity in human (Bertram et al. 2017a; Bertram et al. 2017b) and yeast (*S. pombe* and *S. cerevisiae*; Yan et al. 2015; Wan et al. 2016; Yan et al. 2016; Yan et al. 2017; Bai et al. 2018; Zhan et al. 2018; Wan et al. 2019) spliceosomes.

Multiple similarities in mechanism and certain sequences between spliceosomes and Group II introns, transposable elements that catalyze excision and ligation (Roscigno et al. 1993; Sontheimer et al. 1999; reviewed by Papasaikas and Valcarcel 2016), provided the first evidence that the spliceosomal reaction is also RNA-catalyzed. Crystal structures of the Group IIC intron of *O. Iheyensis* elucidated a triple helical catalytic core involving a bulge in Domain (D)V, the catalytic triad AGC at the base of DV, and a short single stranded junction between DII and DIII to position two Mg^2+^ that promote catalysis (J2/3; Toor et al. 2008), thus providing a clear role of these components in formation of the metal ion-dependent catalytic center of this splicing machinery.

By comparison, the catalytic center of the spliceosome is a complex defined by the U2 and U6 snRNA (Fabrizio and Abelson 1990), the most highly conserved among the five snRNA sequences. Inter- and intra-molecular base pairing between U2 and U6 snRNA lead to the formation of two intermolecular helices, Helix I and Helix II, and the U6 intramolecular stem loop (ISL). The U6 snRNA ACAGAGA sequence of Helix I is involved in 5′ splice site selection, and in promoting both steps of the splicing reaction (Luukkonen and Seraphin 1998; Mefford and Staley 2009), and the U2 snRNA sequence opposing this ACAGAGA segment pairs with a region of the intron to position the branch site residue. Catalytically essential metal ion-binding sites have been identified at the catalytic AGC triad at the base of the U6 ISL, a bulged uridine in the ISL, and nucleotides at the 3′ end of the ACAGA<u>GA</u> loop (Gordon et al. 2000; Yu et al. 1995; Yuan et al. 2007; Yean et al. 2000; Sontheimer et al. 1997).

Analogy between sequence elements and ion-binding sites critical to forming the long-distance interactions and in catalyzing cleavage in the Group II intron (Toor et al. 2008) led to the hypothesis of an analogous motif in spliceosome involving the bulged U of the ISL, AGC triad, and the 3′ end of the ACAGA<u>GA</u> loop (Madhani and Guthrie 1994; Keating et al. 2010). Such an interaction was evidenced by cross-linking and genetic mutations in yeast spliceosomes in their cellular environment (Fica et al. 2014). Cryo-EM based models fully support this model (Anokhina et al. 2013; Yan et al. 2016), and establish the absence of proteins in the immediate vicinity of the catalytic site contact.

The demonstration that a protein-free construct representing the U2-U6 snRNA complex catalyzes a Mg^2+^-dependent branching reaction (Valadkhan et al. 2009) provides evidence that the RNA alone is capable of catalysis. However, very low yield and very slow rate of the modified splicing reaction, as well as the finding that the components forming the active site are far apart in the protein-free state of the yeast (Burke et al. 2012; Guo et al. 2009) and human (Karunatilaka and Rueda 2014) U2-U6 snRNA complexes, support the requirement for spliceosomal proteins to facilitate formation and/or stabilization of an active conformation.

The role of the central junction in positioning of the catalytically essential helices of the U2-U6 snRNA complex has been investigated in yeast and human systems. Although a three-helix conformer has been identified in intact spliceosomes (Yan et al. 2016; Bertram et al. 2017b) and cellular systems (Anokhina et al. 2013), *in vitro* studies of the U2-U6 snRNA complex in its protein-free state have supported formation of two different secondary conformations characterized by three and a four helices about the central junction (Sashital et al. 2004; Guo et al. 2009; Burke et al. 2012; Zhao et al. 2013; Karunatilaka and Rueda 2014) and in dynamic equilibrium with each other (Zhao et al. 2013). The observation of alternative distributions is interesting because the conserved invariant AGC triad is paired with U6 snRNA to extend the base of ISL in the four-helix variant and is paired with U2 snRNA in the three-helix conformation; and may thus relate to the ability to participate in formation of the active site (Sashital et al. 2004). In support of a role for a four-helix conformer, genetic experiments had found that four-helix fold maintains a critical role in an unspecified stage in human cells (Wu and Manley 1989).

Interaction with Mg^2+^ is a necessary and sufficient to facilitate folding of the Group II intron into a catalytic form in the absence of proteins (rev. by Pyle 2002; Kruschel and Sigel 2008), but its role in the folding process of the spliceosomal U2-U6 snRNA complex, particularly in junction conformation and the positioning of helices around the junction, is not clear. In addition to its essential role in forming the catalytic two-metal ion center that catalyzes cleavage and ligation of scissile phosphates of splice sites, we hypothesized that Mg^2+^ maintains important roles in the nonspecific screening of RNA backbones near the central junction that facilitates positioning and flexibility of the complex in the absence of protein components.

To enhance our understanding of the role of the central junction and of divalent metal ions in facilitating formation of the active structure, we have investigated the orientation of the helical stems of the human U2-U6 snRNA complex in the absence of spliceosomal proteins. Toward this goal, we used time-resolved Förster resonance energy transfer (trFRET) to measure distances between stems of the wild type and mutant complexes, and dependence on added Mg^2+^.

Ensemble and single-molecule (sm)FRET techniques have been used to analyze orientations of stems around Holliday junctions in DNA (Clegg et al. 1994), junctions in hairpin and other ribozymes (Walter et al. 1998; Liu et al. 2007) (Tan et al. 2003), Group II introns (Steiner et al. 2008), and spliceosomal junctions (Yuan et al. 2007; Guo et al. 2009; Karunatilaka and Rueda 2014) and time-resolved luminescence resonance energy transfer was used to estimate locations of multiple ion-binding sites in a protein-free U2-U6 snRNA complex (Yuan et al. 2007). However, to the best of our knowledge, this is the first application of trFRET to probe three-dimensional orientations of stems around three- and four-helix junctions, and Mg^2+^-dependent changes in the distribution and conformations of those junctions, in such a system.

Our results showed that the four-helix conformer adopts a roughly tetrahedral arrangement and that the angles between helical stems around the junction decrease upon addition of Mg^2+^, resulting in greater compaction. In contrast, the three-helix conformer associated with intact spliceosomes adopts a closer arrangement in the absence of Mg^2+^, but the stems bearing the catalytically essential elements are markedly distanced from each other upon addition of Mg^2+^. Although the data are consistent with the central junction maintaining a Mg^2+^-sensitive role in positioning of helices, in neither protein-free conformer, in the presence or absence of Mg^2+^, are the regions bearing the catalytically essentially metal ions in sufficiently close proximity to form the interactions associated with the catalytic center.

## RESULTS

To analyze role of the central junction in positioning the two helices critical to forming the active sites in both conformers, and to build a three-dimensional structural model of the orientation of the four helical stems of the majority form of the protein-free human U2-U6 snRNA complex, we used steady state and time-resolved FRET techniques to measure distances between termini of complexes formed by paired fragments representing U2 and U6 snRNA complex. RNA constructs are described in Materials and Methods and shown in Figure 1.

**Figure 1:**
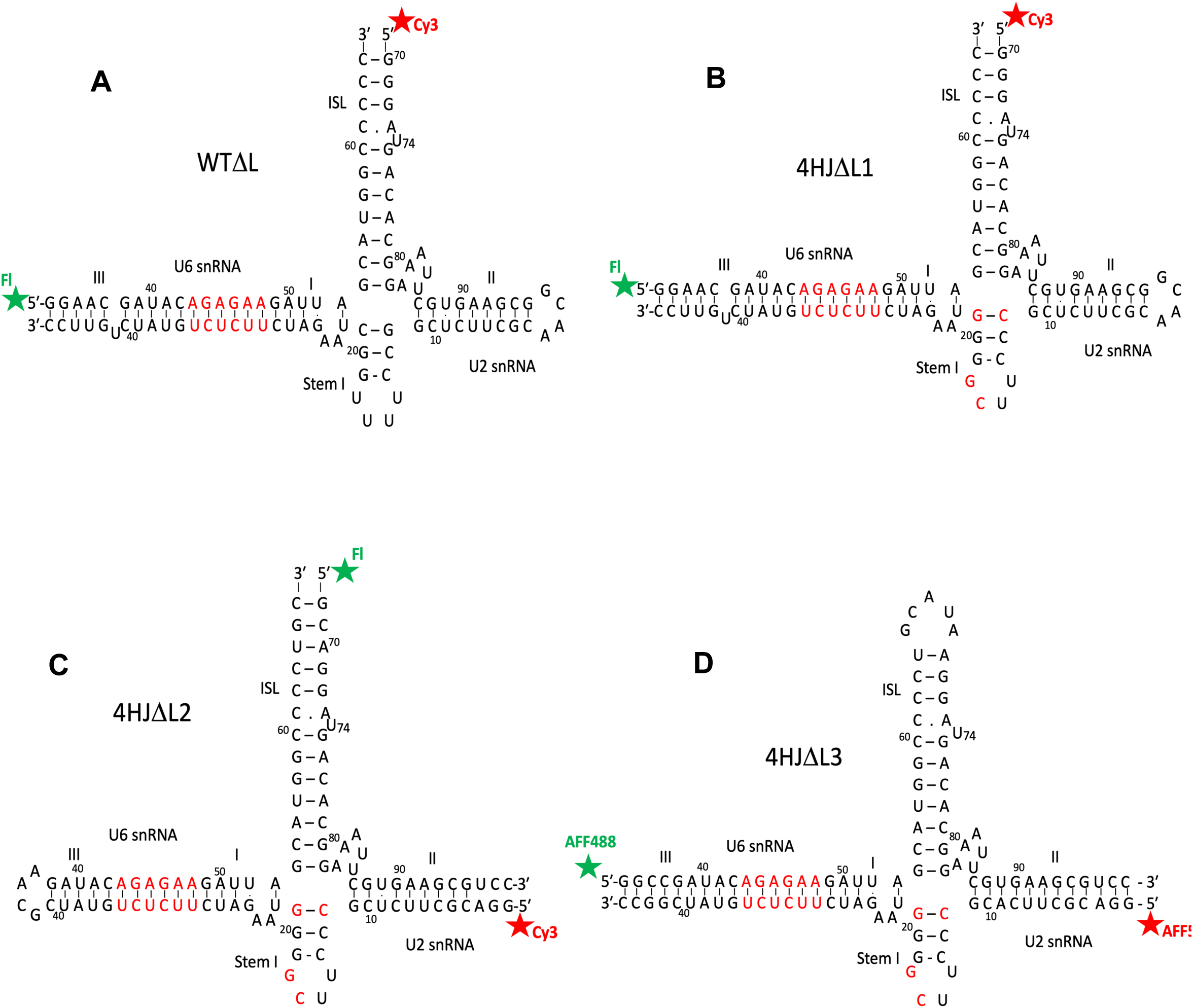
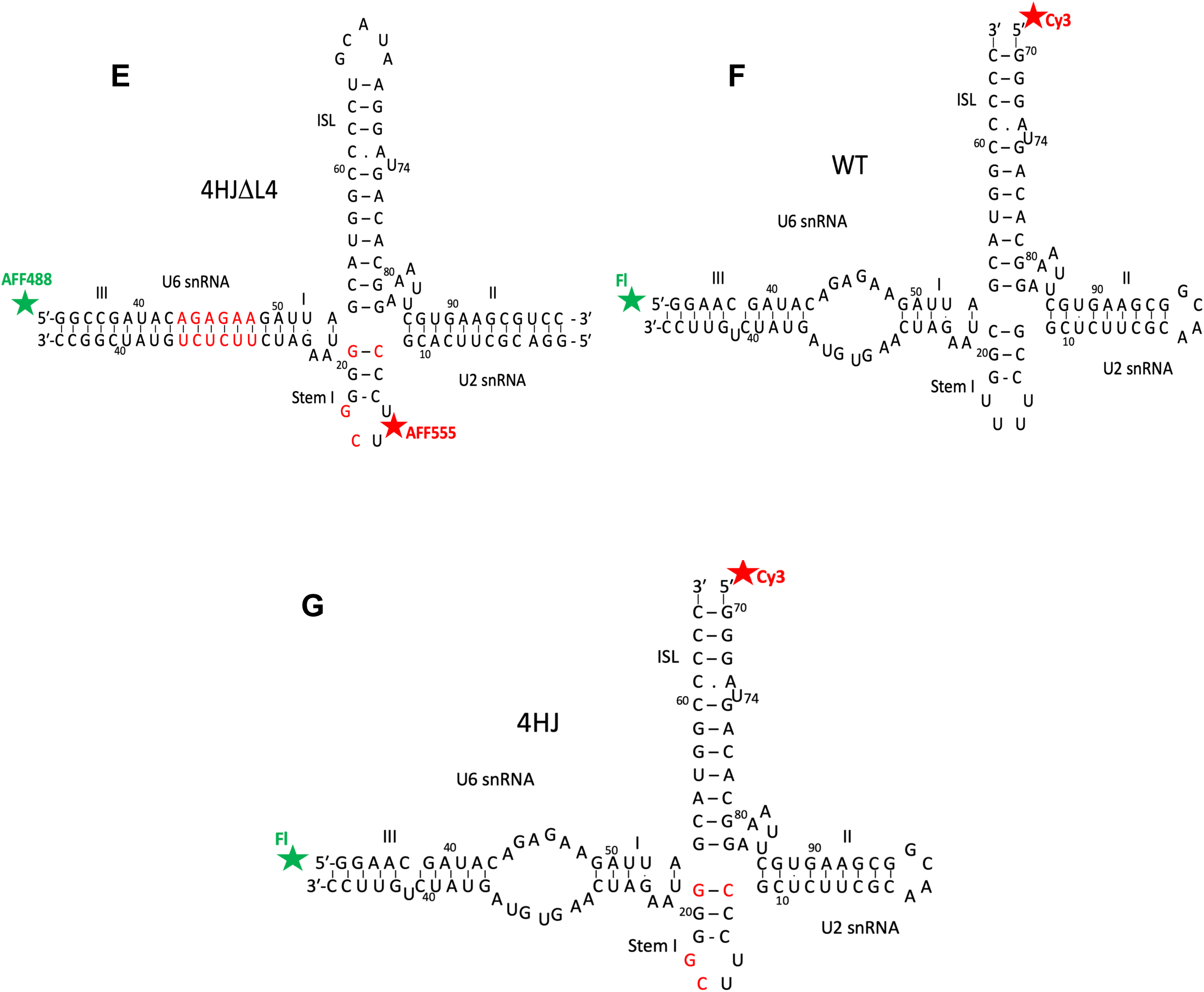
Constructs representing the human U2-U6 snRNA complex used in these experiments. Modifications to helix length including truncation, deletion or addition of hairpin loops to create “chimeric” strands, or nucleotide changes to favor base pairings, are described in Materials & Methods. Constructs are all depicted in the four-helix secondary structure; the AGC catalytic triad corresponds to nucleotides A_53_G_54_C_55_ of U6 snRNA. (*A*): Wild-type junction in which the ACAGAGA loop is “closed” by pairing with a complementary sequence (shown in red), with Fluorescein as the Donor (D; Fl) and Cy3 as the Acceptor (A) on termini of helix III and U6 ISL, respectively (WTΔL); (*B*): mutations in Stem I favor the four-helix conformer (individual nucleotide changed in red), also with a “closed” ACAGAGA loop and dyes on helix III and U6 ISL (4HJΔL1); (*C*): four-helix mutant with “closed” ACAGAGA loop, and Fl and Cy3 dyes on termini of U6 ISL and Helix II, respectively (4HJΔL2); (*D*): four-helix mutant with “closed” ACAGAGA loop with AF488 (D) and AF555 (A) dyes on termini of Helix III and Helix II, respectively (4HJΔL3); (*E*): four-helix mutant with “closed” ACAGAGA loop with AF488 and AF555 dyes on Helix III and Stem Loop I, respectively (4HJΔL4); (*F*): wild-type U2-U6 junction with wild type ACAGAGA loop, and Fl and Cy3 on termini of Helix III and U6 ISL, respectively; (*G*): four-helix mutant with ACAGAGA loop, with Fl and Cy3 dyes on termini of Helix III and U6 ISL, respectively (4HJ).

**Figure 2:**
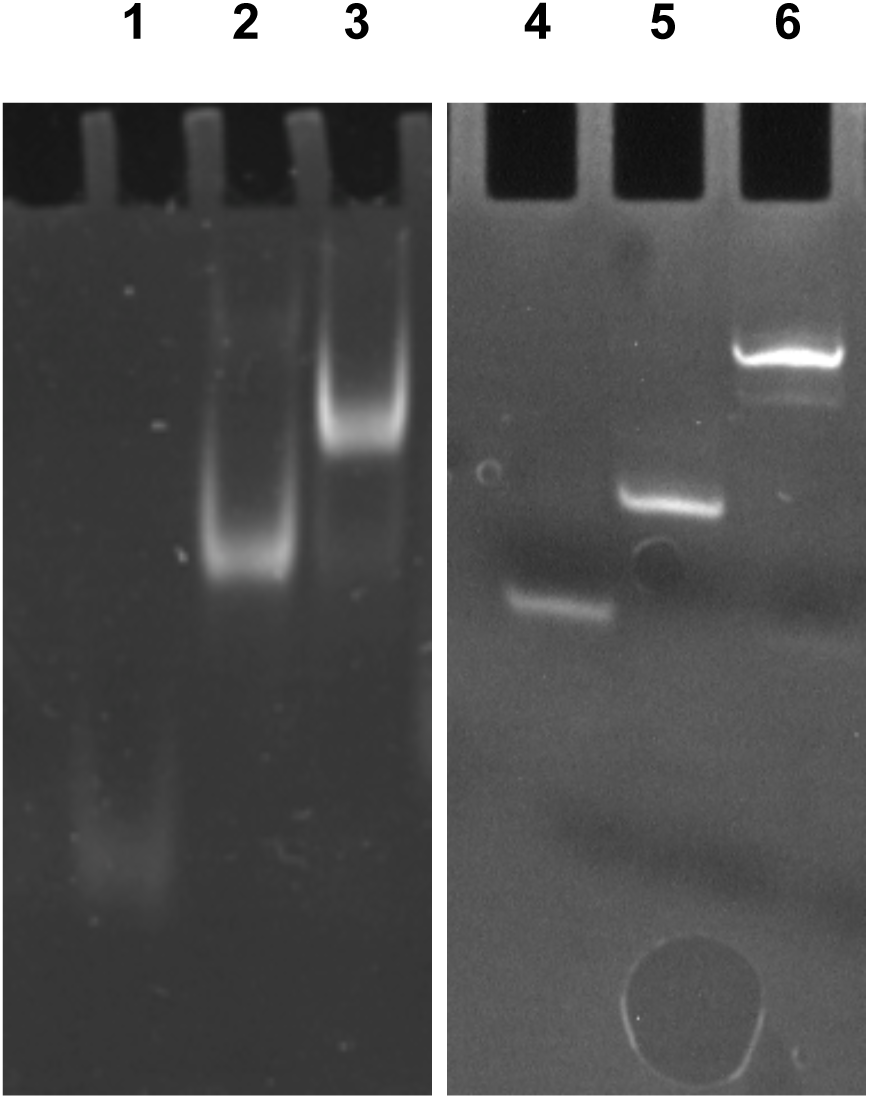
Non-denaturing polyacrylamide gel electrophoresis used to analyze pairing of equimolar concentrations of RNA strands of U2-U6 snRNA constructs. **Left:** Pairing of strands of construct F (see legend of Figure 1F). Lane 1: 32 nucleotide (nt) fragment representing the 5′ region of U6 snRNA (Helix III through the 5′ side of the U6 ISL); Lane 2: 74 nt U6-U2 “chimeric” fragment containing the 3′ side of U6 ISL linked to the U2 strand *via* a GCAA tetraloop; Lane 3: Equimolar concentrations of strands in Lane 1 and Lane 2 paired to form construct F. The shift shown in Lane 3, with no residual band in Lanes 1 or 2, indicates complete and stoichiometric pairing between strands. **Right:** Pairing of strands of construct D (Figure 1D). Lane 4: 44 nt U2 snRNA strand; Lane 5: 65 nt U6 snRNA strand; Lane 6: Equimolar concentrations of strands in Lane 4 and Lane 5 paired to form construct D. The shift shown in Lane 6, with no residual band in Lanes 4 or 5, indicates complete and stoichiometric pairing between strands.

### Steady state FRET measurement of the human U2-U6 snRNA complex

We first acquired inter-dye distance measurements between termini of Helix III and the U6 ISL of a U2-U6 snRNA construct, the two stems containing elements that interact to define the catalytic center. This construct (WTΔL; Figure 1A) replaced the sequence of U2 snRNA opposing the ACAGAGA loop with a complementary duplex to create a continuous stem. Although nucleotides in the 5′ end of the U6 and the 3′ end of the U2 snRNA sequences are capable of forming nine Watson-Crick base pairs to form Helix III, this duplex is not observed in cellular (Anokhina et al. 2013) or spliceosomal (Yan et al. 2016; Bertram et al. 2017a) systems. However, this helix forms in the protein-free RNA complex and is useful in these experiments for stabilizing the position of the attached dye on the 5′ terminus of the U6 snRNA fragment. In this case we measured FRET between a fluorescein Donor (D)-labeled 5′ end of the abbreviated U6 snRNA strand and the Cy3 Acceptor (A)-labeled U6 ISL.

Steady state measurement of FRET efficiency between D and A, calculated from the decrease in emission of D in the presence of A (corrected for emission of A alone when illuminated at the excitation wavelength of D, 495 nm; see Materials and Methods), was 13.7% (Figure 3). Using equation 4 (Materials and Methods) and R_0_ = 56 Å for the fluorescein-Cy3 dye pair, this transfer efficiency corresponds to a mean distance of ∼76 Å between the dyes. We note that previous enzymatic structure probing and ^19^F NMR experiments from our laboratory identified conformational heterogeneity of the central junction of the protein-free human U2-U6 snRNA complex characterized by a mixture of three- and four helix conformers (Zhao et al. 2013) and identified the predominant fold in the protein-free human U2-U6 snRNA complex under similar conditions as used here as the four-helix conformer. However, spectra derived from steady state ensemble FRET measurements do not provide information to distinguish individual distances in a heterogeneous system.

**Figure 3:**
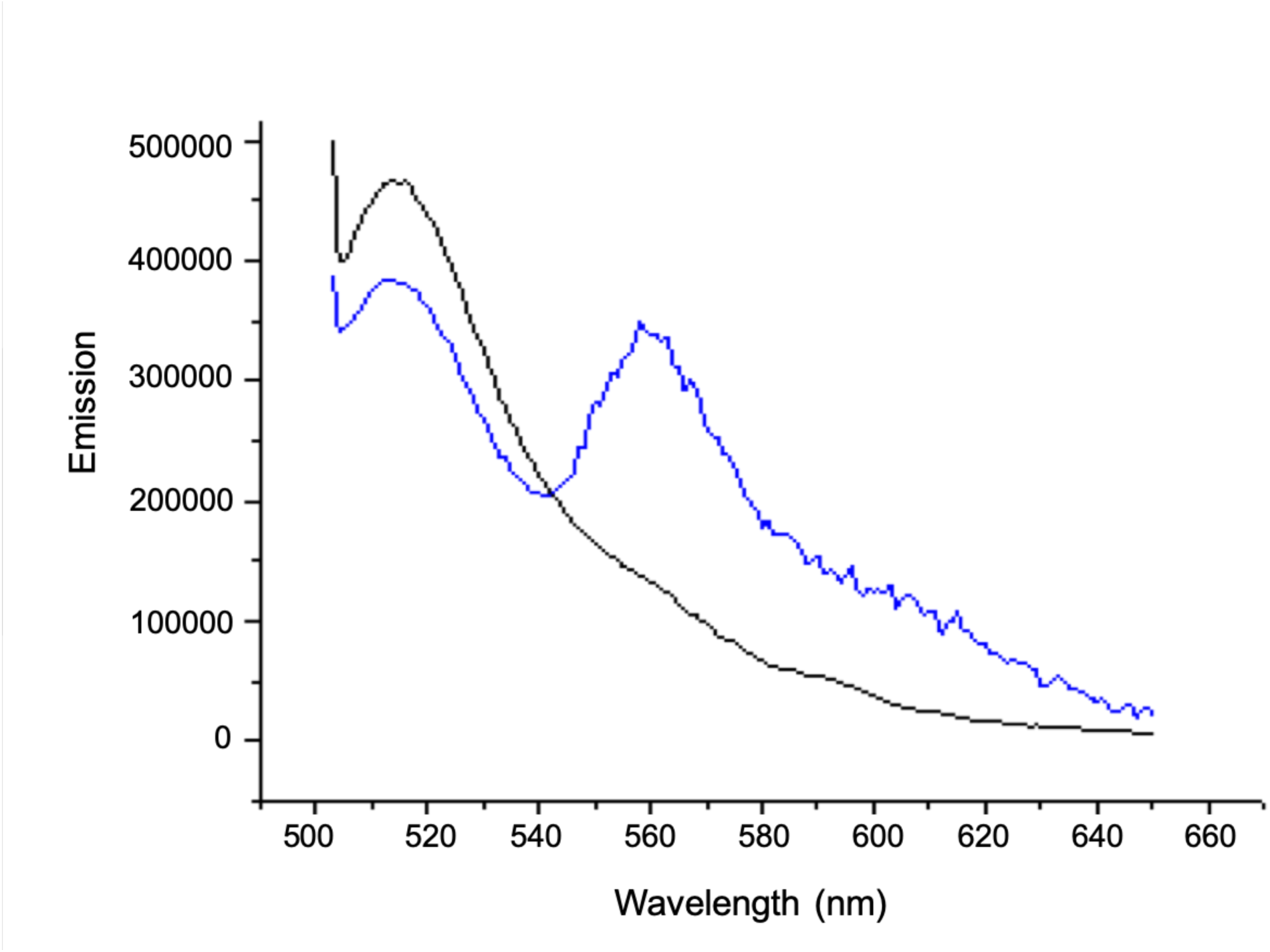
Steady state emission spectrum of fluorescein (Donor) attached to the 5’ terminus of U6 snRNA as the only dye (black line), and with Cy3 labeled at 5’ of U6-U2 chimeric RNA (blue line) in construct WTΔL (Figure 1A) following excitation at 495 nm. Samples were 150 nM RNA in Tris (30 mM)-HEPES (60 mM), pH 7.6, 30 mM NaCl, 1 mM EDTA (no added Mg^2+^). The emission of an RNA sample with Cy3 (the acceptor) only following excitation at 495 nm (not shown) was subtracted from the FRET trace to eliminate contribution of direct excitation of Cy3 at this wavelength.

### trFRET of a U2-U6 snRNA complex with the wild type junction

To estimate distances in each of the major conformers in this heterogeneous system, as well as the distribution of the populations, we utilized trFRET. The advantage of this approach is that the decay curve of a pulse-excited donor dye transferring energy to a suitable acceptor dye contains rate/distance information for all D-A distances represented in an ensemble weighted according to fractional representation in the ensemble; deconvolution of decay curves thus allows determination of the relative populations of signals from multiple conformers in a heterogeneous system.

Using the same WTΔL construct, we first measured the decay of emission of D alone (with an unlabeled U6-U2 chimeric strand), followed by D in the presence of A, for Construct WTΔL (Figure 1A), and fitted decay curves with single exponential or the weighted sum of two exponential curves (Figure 4). The time-resolved decay curve of the D alone was fitted by a single exponential curve with a lifetime,τ, of 4.05±0.008 ns and a CHI-square (goodness of fit; χ^2^) value = 1.07. The decay curve for D in the presence of A was first fit with a single exponential curve with τ = 3.69±0.010 ns and χ^2^ = 2.67, *i.e.* a much poorer fit. Fitting the curve with two decay components, however, with lifetimes of 3.87±0.01 ns (fractional amplitude of 91.4%) and 0.89±0.06 ns (fractional amplitude of 8.6%), for distances between the dyes of 93.0 Å and 45.3 Å, respectively, yielded a far better overall fit (χ^2^ = 1.15). These two lifetimes correspond to an average distance of 80.0±2.2Å, similar to the result from steady state ensemble FRET. Our earlier finding that the four-helix conformer was by far the larger fraction (Zhao et al. 2013) suggests that it is likely to correspond to the larger distance, and that the minor three-helix conformer is associated with the shorter distance. The next task was to match each set of distances with the conformer it represents independently without any assumptions.

**Figure 4:**
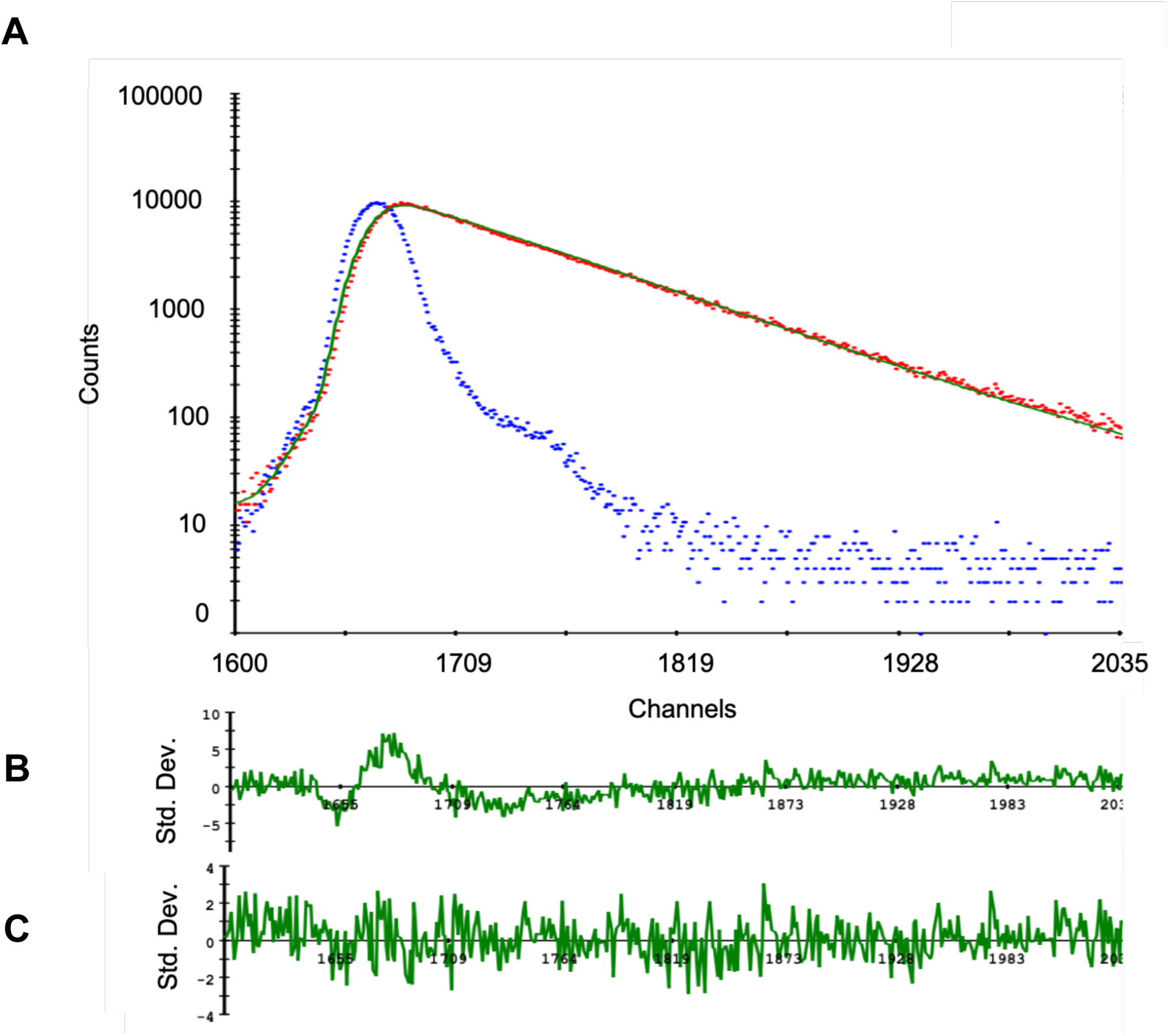
The exponential decay fitting process for (*A*): fluorescein (Donor) undergoing FRET to Cy3 (Acceptor) in Construct WTΔL. The blue decay curve is instrument response. The red decay curve is for fluorescein in response to pulses at 495 nm. The solid green curve was a best fit by a biexponential curve corresponding to two lifetimes of 3.87±0.01 (91.4%) and 0.89±0.06 ns (8.6%). (*B*) Display of residuals derived from fit with a mono-exponential curve indicates a poor fit (CHISQ=2.67); (*C*) Display of residuals derived from fit with a biexponential curve indicates a better fit (χ^2^ =1.15).

### trFRET of the four-helix conformer

To obtain distances related to individual conformers, we performed measurements using construct with a modification of nucleotides near the junction that favored the four-helix structure (construct 4HΔL1; Figure 1B) and compared results with those for the wild type junction (WTΔL). Measurements of trFRET between the dyes in the construct 4HΔL1 labeled at the same sites as of construct WTΔL indicated a fitted single exponential curve with a distance of 90.8±0.6 Å (χ^2^ = 1.24), a value very similar to that of the deconvoluted four-helix conformer and very different from that of the minor conformer, confirming that the longer lifetime component in the decay curve deconvolution corresponds to the four-helix conformer, and the smaller component with the shorter lifetime therefore corresponds to a three-helix conformer.

### Translation of distance information into junction angles

From the distance between termini of helical stems and estimated lengths of the stems (Helices III-I and ISL), we estimated angles between stems. For construct 4HΔL1, the stem formed by Helices I and III is a continuous A-type helix of 24 nucleotides (2.6 Å between centers of base pairs); with 8.5 Å extension for fluorescein and a 2 carbon amino linker and 5 Å due to addition of four more C-C bonds in six-carbon amino linker compared to two-carbon amino linker, the total length is ∼(2.6 Å x 24) + 8.5 + 5 =75.9 Å. The acceptor-Cy3 attached at U6 ISL stacks onto the RNA helix and therefore contributes to the length of the human U6 ISL as one additional nucleotide, for a total of 33.8 Å. Triangulation of the lengths of the two stems and the distance measured by FRET in the heterogeneous wild type junction produced an average angle of 84.4±3.6°. We note that this value represents a mean between two conformers with very different distances, and therefore of limited value. Calculated from individual distances obtained from deconvolution of the WT decay curve, 93 Å and 45.3 Å, we obtained values of ∼109.9° for the angle in the majority four-helix conformer, *i.e.* a large distance between the ISL and Helix I/III stems (Figure 6A); calculation of the same angle in the 4HΔL1 construct from the directly measured distance was 105.2±1.2°, very similar to that obtained by deconvolution. In contrast, a distance of 45.3 Å in the three-helix conformer corresponds to a much smaller angle of 19° (Figure 6C), indicating that even in the absence of all sequence elements necessary for formation of the catalytic triplex interaction (pairing of the ACAGAGA loop prevents any tertiary interaction), the three-way junction positions the ISL and Helix I/III stems in a much closer orientation than it does in its four-helix conformation.

We also estimated the distance between U6 ISL and Helix I/III values for the minority three-helix conformer by subtraction of the measured decay curve for the fraction of four-helix fold (91.4%) from that of the (heterogeneous) WT. The decay curve of four-helix fold is well fitted to a single exponential decay curve with correlation coefficient, R^2^, = 0.9999, whereas the subtracted decay curve for three-helix conformer is less perfectly fitted to a single exponential decay curve shown in Figure 5A (R^2^=0.97, calculated using SigmaPlot 13.0; Systat Software, Inc., San Jose California USA; www.systatsoftware.com), perhaps as the result of several sub-conformers; a similar conclusion was reached from marked ^19^F peak heterogeneity in NMR experiments (Zhao et al. 2013). The distance calculated between U6 ISL and Helix III for the remaining three-helix is 56.3 Å, corresponding to an angle of 43.3°. Although there is a noticeable difference in the angle obtained by deconvolution and subtraction (19° vs 43.3° respectively), this disagreement may result from uncertainty associated with dye/linker mobility, heterogeneity of three-helix conformer, the large difference in helix length, and/or the indirect nature of the calculation involved for analysis of this minority conformer. However, both values indicate an acute angle in the range consistent with the Y shape structure modeled by Butcher and coworkers shown previously for the yeast U2-U6 snRNA complex (Burke et al. 2012), implying that the junction has a role in positioning the two helices near to each other.

**Figure 5:**
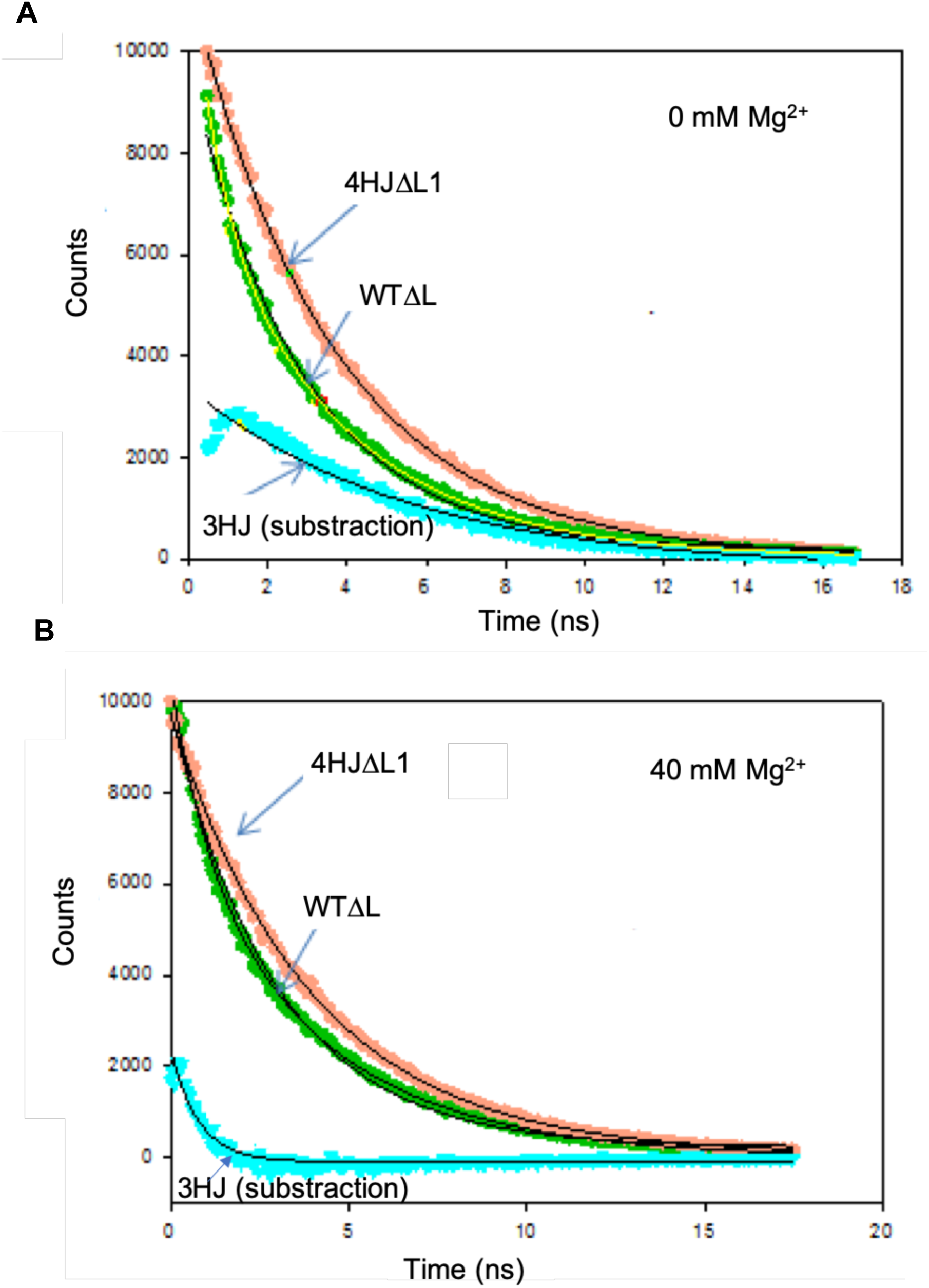
Calculation of a decay curve for the three-helix conformer (cyan) by subtraction of the experimentally obtained decay curves of fluorescein-labeled 4HJΔL1 (orange) from that of WTΔL (green). (*A*) Curves without added Mg^2+^ and (*B*) curves with added Mg^2+^ (40mM). Concentration of dye-labeled RNA was 150 nM. trFRET measurements were recorded on a timescale of 20 ns to a total of 10,000 counts in the peak. The decay curve of fluorescein on 4HJΔL1 and WTΔL were fit by a mono-exponential and biexponential curve, respectively, with an excellent R^2^ (R^2^>0.999). The calculated decay curve for the three-helix conformer in each case was fitted with a mono-exponential curve yielding a less perfect fit (R^2^ value ∼0.974 and 0.911 without and with Mg^2+^, respectively), suggesting a mix of several three-helix conformers (see Results for details).

### Measurement of other distances in the four-helix conformer

To contribute to the three-dimensional topology models of the two conformers of the protein-free human U2-U6 snRNA complex, we also determined angles between dye pairs at other sites within the complex. In each case, we repeated trFRET measurements to determine inter-dye distances using constructs with dyes on different termini, and with either the wild type or the helix mutated to form only the four-helix conformer. As in WTΔL and 4HΔL, the ACAGAGA loop was eliminated by extensive WC pairing.

To measure the distance between the cleaved hairpin loop of U6 ISL and the terminus of Helix II we created a four-helix construct, 4HΔL2 (Figure 1C), and labeled 5′ termini with fluorescein and Cy3, respectively. Results of trFRET measurements indicated a distance of 73.0±1.0 Å, triangulated into an angle of 110.1±2.4°. This angle calculation included an estimation that five U6 snRNA unpaired nucleotides (AAAUU) at the junction form a loop contributing ∼5 Å to the length of Helix II.

For measurements between Helices II and I/III, we labeled the 5′ termini of strands representing U2 snRNA (in Helix II) and U6 snRNA (in Helix III) (4HJΔL3; Figure 1E). We used the AlexaFluor dye pair (AF488 and AF555) to take advantage of the longer R_0_ for the larger anticipated distance between the termini (Yuan et al. 2007). AF488 and AF555 have been shown by simulation to be relatively mobile in aqueous solution (Corry and Jayatilaka 2008); we therefore assumed they would behave like fluorescein in calculation of the length of RNA duplex. The distance between probes at 5′ termini of U2 and U6 snRNA was 97.0± 1.0 Å, which triangulated to an angle of 116.7±2.0°.

Finally, to measure the distance between Stem I of U2 snRNA and Helix III (4HJΔL4; Figure 1E), we measured trFRET between AF488 labeled at 5′ of U6 snRNA and AF555 at internally labeled U_16_ of U2 Stem I. The distance from 5′ U6 snRNA to U2 stem I is 85.2±0.8 Å, for an angle of 139.0±4.0°. These data have allowed us, for the first time, to create a three-dimensional model of the human protein-free U2-U6 snRNA complex in its four-helix form (Figure 6A) and calculated three-helix form (Figure 6C).

### Effect of Mg^2+^ on conformation of the human U2-U6 snRNA complex

Both spliceosomes and Group II introns are dependent upon interaction with Mg^2+^ for assembly, including interaction of key sequence elements, as well as for catalytic activity. To investigate the impact of Mg^2+^ addition on the three-dimensional fold of the protein-free human U2-U6 snRNA complex, we repeated each of the measurements of interhelical distances in the presence of MgCl_2_ from concentrations of 2 mM (near cellular concentration; Romani and Scarpa 1992) to 40 mM (the value used in previous *in vitro* studies; Guo et al. 2009; Zhao et al. 2013). Results indicated a gradual decrease in distance between termini of Helix I/III and U6 ISL in the four-helix conformation (4HJΔL1) with each addition (Figure 7). At a final concentration of 40mM Mg^2+^, the distance decreased from 90.8±0.6Å to 78.3±0.8 Å, for a decrease in angle from 105.2±1.2° to 81.3±1.4°, *i.e.* a closer proximity of Helix I/III and U6 ISL. The distance between Helix II and U6 ISL in construct 4HJΔL2 (Figure 1C) decreased by 5.1Å, for a decrease in angle from 110.1±2.4° to 99.1±2.0°. No significant change was observed for distance between Helix II and Helix I/III (in 4HJΔL3). The distance between Stem I and Helix I/III in Construct 4HJΔL4 was lengthened slightly by 2.1Å for an increase in angle from 139.0±4.0° to 150.5±5.6°, placing Stem I further “behind” the junction (Figure 6B). All changes in the four-helix conformer upon addition of Mg^2+^ appear to be in the approach of helices toward the U6 ISL, originating from changes in stem orientation (Figure 6B), resulting in a more compact folded structure. We had previously noted this compaction by increased migration measured by analytical ultracentrifugation in the presence of 40mM Mg^2+^ (Zhao et al. 2013).

**Figure 6:**
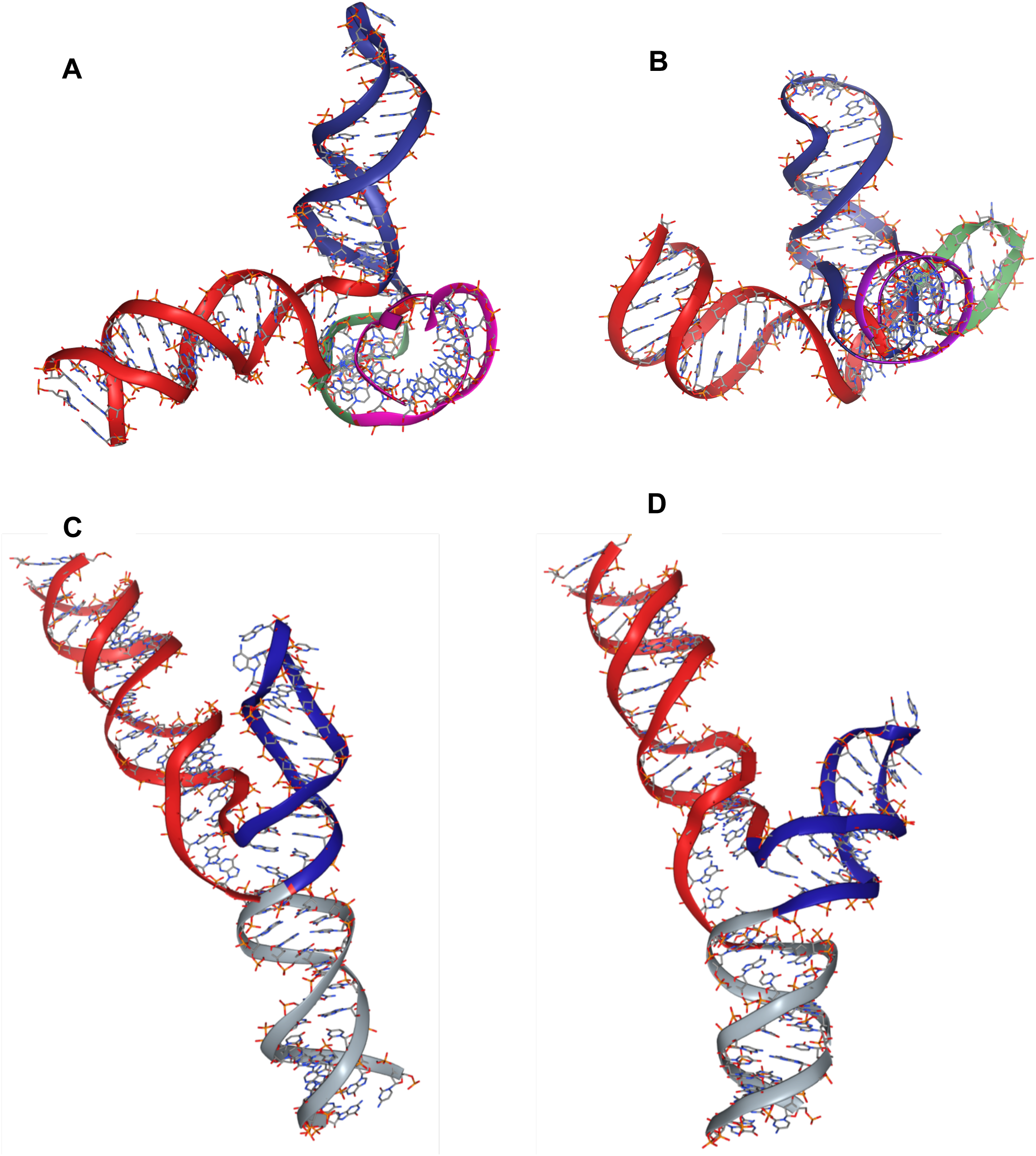
Predictive computational models of the two major conformers of the human U2-U6 snRNA complex obtained by simulation using SimRNA (Magnus et al. 2016), using distance restraints between dyes determined experimentally and visualized by NGLview (Nguyen et al. 2018). (*A*) and (*B*): Models of the four-helix junction conformer (Helix I/III in red; U6 ISL in blue; U2 Stem I folded behind the junction in green; Helix II in violet) (*A*) without Mg^2+^ and (*B*) with 40 mM Mg^2+^, showing a compaction of Helix I/III, ISL, and Helix II around the junction (Stem I is folded behind the junction in this view). (*C*) and (*D*): Models of calculated three-helix junction conformer with three helices (Helix I/III in red; U6 ISL in blue; Helix II in grey) (*C*) without Mg^2+^ and (*D*) with 40 mM Mg^2+^, showing a widened junction. The position of Helix II in the three-helix conformer was not experimentally determined but found to be in the lowest energy position by simulation.

**Figure 7:**
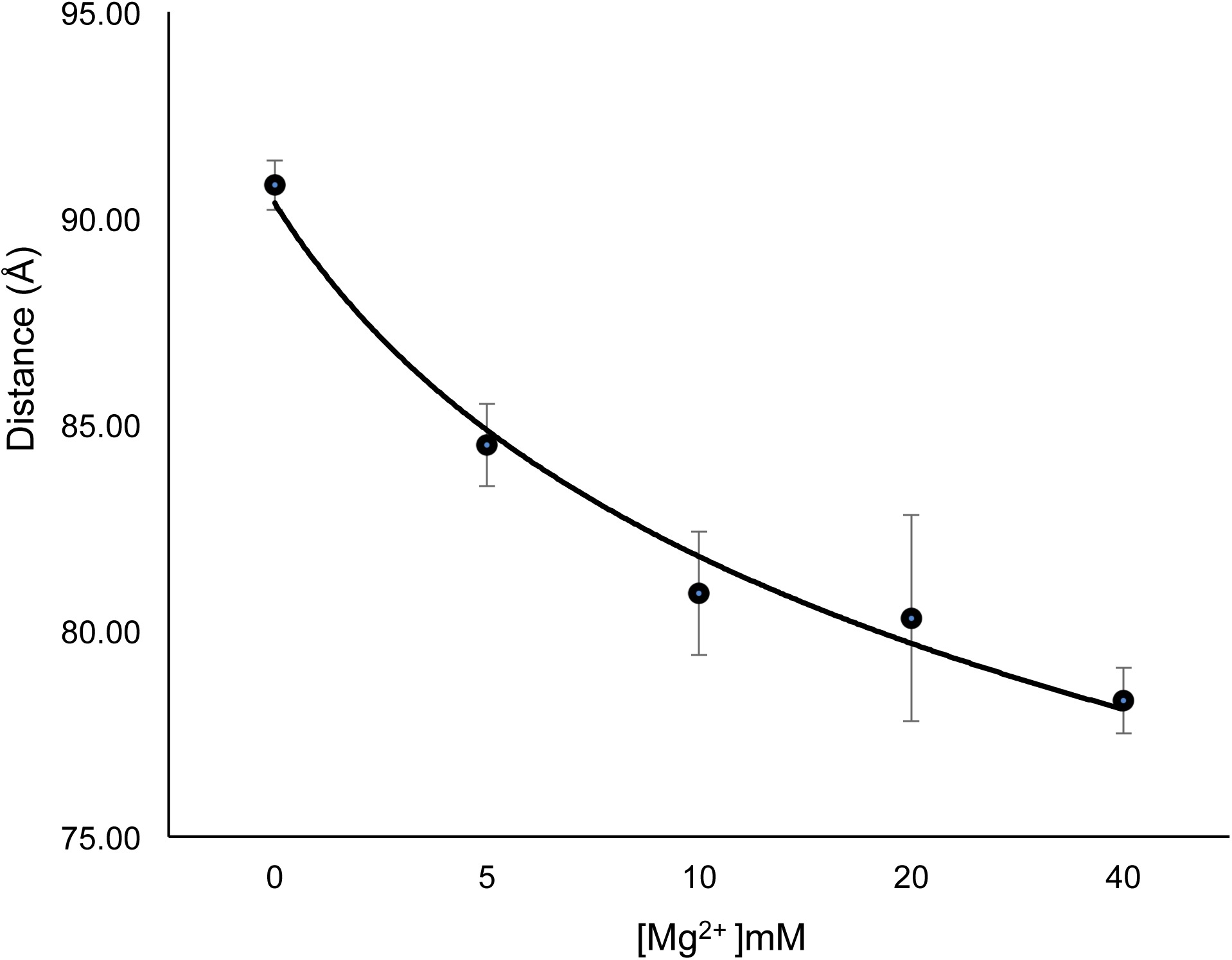
Decrease in distance between dyes attached to Helix I/III and U6 ISL in construct 4HJΔL1 with respect to increasing Mg^2+^ concentration from 0 to 40 mM. Error bars indicate ± standard deviation. Experimental details in Materials and Methods.

We then investigated changes in the distribution of four- and three-helix conformers upon addition of 40 mM Mg^2+^. The mean distance between Helix I/III and U6 ISL in heterogeneous WTΔL shortened by 3.5 Å with added Mg^2+^, a value much smaller than the change in four-helix construct (12.5 Å). However, deconvolution of the decay curves of WTΔL upon addition of Mg^2+^ revealed approximately twice the fraction of the three-helix conformer relative to its fraction in the absence of Mg^2+^ (increase from 8.6% to 17% of the total), but also induced marked changes in stem orientation. By deconvolution, the distance between Helix I/III and U6 ISL in the three-helix junction conformer in 40mM Mg^2+^ increased to 67.7 Å (from 45.3Å), triangulated to an angle of 63.1° (from 19°); calculation by subtraction (Figure 5B) yielded a change in distance to 68.9 Å (from 56.3 Å) triangulated to an angle of 65.2° (from 43.3°). By either approach, this change agrees with data from smFRET studies illustrating that addition of Mg^2+^ widens the distance between these stems in the three-helix conformer (Guo et al. 2009). Thus, the effect of Mg^2+^ on the three-helix conformer (Figure 6D) is opposite to what we observed in four-helix conformer (Figure 6B).

### Role of ACAGAGA loop in helix orientation

To examine the effect of the ACAGAGA loop (*i.e.* an open loop in the presence of the naturally occurring U2 snRNA sequence) to the folding of the complex, we performed the same measurements for the distance between Helix I/III and U6 ISL with the wild-type complex (Construct WT; Figure 1F) and the four-helix mutant (Construct 4HJ; Figure 1G), both with a native (open) ACAGAGA loop. In both cases, results of trFRET measurements show a small increase of inter-dye distances, from 80.0±2.2 Å (construct WTΔL) to 84.0± 1.5 Å (construct WT); and from 84.0±1.2 Å (4HJ) to 90.8±0.6 Å (4HJΔL1), suggesting that the ACAGAGA loop contributes a small kink to Helix I/III in the protein-free U2-U6 snRNA complex. There was also no observable change in measurements repeated on constructs in which the region of U2 snRNA was paired with a short intron fragment representing the branch site (pairing confirmed by change in electrophoretic mobility of components using nondenaturing PAGE; data not shown). We had noted previously that the distribution between junction conformations was not altered significantly in ^19^F-NMR spectra in response to the presence or absence of the ACAGAGA loop (Zhao et al. 2013). In none of these cases did inclusion of the native ACAGAGA loop, with or without pairing to an intron segment, facilitate increased proximity or interaction between Helix I/III and the U6 ISL.

## DISCUSSION

Our goal was to analyze the contribution of the central junction of the spliceosomal U2-U6 snRNA complex in facilitating splicing activity by concerted positioning of the helices associated with formation of the catalytic center. Using protein-free human snRNA complexes labeled at termini or internally with fluorescent dyes, results acquired from trFRET measurements enabled characterization three-dimensional arrangements of helical stems about the central junction of the human U2-U6 snRNA for the two major folds in the absence of proteins, and changes upon incubation of Mg^2+^.

The spliceosome’s U2-U6 snRNA complex is characterized by helical stems radiating from a central junction. Alternative conformers of the junction, characterized by three- and four-helix arrangements of the stems, have been previously documented (Zhao et al., 2013). The U6 snRNA ISL and intramolecular duplex including the U6 ACAGAGA loop and U2 segment that pairs with the intron branch site sequence (Helix I/III in Fig 1) are the same in both junction conformers; the feature that differs between the two is that the AGC catalytic triad (nucleotides 53-55 in constructs shown in Figure 1) in the four-helix junction pairs with the 3′ stem of the U6 ISL, whereas in the more open three-helix conformer, it pairs with U2 snRNA, and thus may be positioned more effectively for interaction with the ISL and ACAGAGA loop to create the active site.

Genetic studies by Manley identified the importance of formation of the four-helix conformer in human cells (Wu and Manley 1989), and solution NMR studies by Butcher and coworkers identified this junction structure in yeast U2-U6 snRNA complexes *in vitro* (Sashital et al. 2004). In contrast, genetic studies in yeast by Madhani and Guthrie (1992) identified a three-helix conformer as the critical conformer. This conformation was supported by single-molecule FRET (Guo et al. 2009), and by a combination of NMR, SAXS, and molecular modeling studies (Burke et al. 2012) of yeast U2-U6 snRNA constructs demonstrated a three-helix junction in which the U6 ISL coaxially stacked with the U2-U6 Helix Ib (Madhani and Guthrie 1992). Interestingly, the U2-U6 snRNA construct used in these latter studies differed only by several base pairs in Helix II, well outside the junction, from that used to identify the four-helix junction by the same group (Sashital et al. 2004). These studies suggest the likelihood that the two alternative conformations with a small energy difference and distribution between the two may depend heavily on experimental conditions. This conclusion is consistent with observations of changes in the distribution of high- and low-FRET in single-molecule studies of U2-U6 snRNA complex of yeast (Guo et al. 2009) and human (Karunatilaka and Rueda 2014) spliceosomes that were attributed to four- and three-helix conformers.

In an attempt to resolve the disparate models, in previous published work we probed the conformation of the protein-free human U2-U6 snRNA complex biochemical structure probing and ^19^F NMR techniques to analyze conformations of the central junction of the protein-free human U2-U6 snRNA (Zhao et al. 2013); the major ^19^F peak corresponding to the four-helix conformer was sharp, suggesting that the four-helix conformer is likely to be a single fold, while the minority very broad and heterogeneous peak associated with single-stranded environment for the fluorinated U within the junction region and indicative of a three-helix conformer in the human U2-U6 snRNA complex, there may be a number of orientations for conformers having three helices. We calculated a very small difference in ΔG between the four-helix and (summated) three-helix conformers from NMR peak volumes (4.7 kcal/mol).

We measured distances for the predominant four-helix fold of the junction by direct analysis of a mutant forming only the four-helix conformer and also by deconvolution of the fluorescence decay curve obtained for the heterogeneous wild type junction. In this first model of the four-helix junction model, we found equivalent results by either method: in the absence of Mg^2+^, the junction adopts a roughly tetrahedral arrangement of helical stems, but upon addition of Mg^2+^, the angle between Helix I/III and the ISL decreases markedly (from 105.2±1.2° to 81.3±1.4°) with increasing concentration of Mg^2+^ (up to 40 mM). This change was accompanied by a decrease in the angle between the ISL and Helix II (change from 110.1±2.4° to 99.1±2.0°). Approach of these two intermolecular stems to the intramolecular (U6) ISL suggests overall compaction of the complex. This finding agrees with a decrease in the Stokes radius and axial ratio (a/b) measured in the presence of Mg^2+^ measured by analytical ultracentrifugation (Zhao et al. 2013).

In contrast, distances between the same helical stems in the minority fraction, characterized by the more open three-helix junction, were determined by deconvolution and also by subtraction of the component corresponding to the four-helix conformer, with qualitatively similar results. Either way, the three-helix conformer exhibited a close position of the stems in the Mg^2+^ free state (angle 19° by deconvolution, 43.3° by subtraction), but responded to Mg^2+^ by a greatly increased separation between the catalytically essential elements (63.1° by deconvolution, 56.3° by subtraction). Moreover, addition of Mg^2+^ to complexes containing the heterogeneous wild type junction displayed a partial conversion from the four- to the three-helix conformer (∼8.6 to ∼17% of the total). Inclusion of the ACAGAGA loop (with or without pairing with an intron fragment to simulate the branch site helix) does not alter these stem orientations measurably. Therefore, the presence of Mg^2+^ induces changes in the equilibrium distribution of junction conformers as well as the positioning of helical stems around the junction.

Our previously published ^19^F-NMR data similarly indicated a shift in the distribution of conformers to increase the relative population of the three-helix conformer from ∼13% of the total to ∼17%, corroborating the fitting of the decay curve obtained by trFRET and further supporting the conclusion that interaction with, or screening by, Mg^2+^ shifts the conformation of the junction toward the AGC presentation that facilitates catalytic site formation. The single molecule FRET studies by Rueda and coworkers on the protein-free yeast U2-U6 snRNA complex showed the coexistence of at least three conformations: four-helix structure, intermediate and three-helix structure (Guo et al. 2009). It is possible that such an intermediate, in equilibrium with other three-helix sub-conformers, contributed to the broad and heterogeneous peak observed in our ^19^F spectra (and would not have been directly detected by either our NMR or ensemble trFRET approaches). If so, our estimate of ∼13% for the population of three-helix sub-conformers is actually an overestimate of the population of several three-helix folds or intermediate(s) and is likely to explain the difference between measurements of distribution by NMR (Zhao et al. 2013) and trFRET (this work). Our NMR acquisition including a maximum of 5 mM Mg^2+^ and ∼0.35 mM RNA (for a ratio of ∼14:1), considerably less than the current ratio of 40 mM Mg^2+^ to 150 nM RNA (267,000:1), so this difference in conditions may contribute to some variation in observed behavior. Not surprising for two conformers with such a small difference in energy between two conformers, we found that the three- and four-helix conformers of the junction exchange on a sub-second time scale (Zhao et al. 2013).

Rueda and coworkers also observed a significant Mg^2+^ dependence in the smFRET data of yeast U2-U6 snRNA complexes labeled at the 5′ terminus of U6 and the U6 ISL loop, noting that the population of complexes displaying high FRET efficiency (assigned to the four-helix conformer) decreased from 64% to 19% (Guo et al. 2009). However, they attributed this change entirely to conversion from four-helix to three-helix junction and did not address the possibility of ion-induced change in helix orientation in either junction conformer. In these experiments, we monitored changes in multiple distances for both the four-helix mutant and wild type (mix of four- and three-helix junctions) U2-U6 snRNA complexes with increasing concentrations of Mg^2+^, from which we calculated change in fraction and orientations of the junction. By either analysis, results of this experiment provided clear evidence that Mg^2+^, in addition to increasing the fraction of three-helix conformer, induces a marked increase in the angle between Helix I/III and the ISL (as well as a more moderate decrease in the angle between these stems in the four-helix conformer). The marked increase in separation between Helix I/III and the ISL upon interaction with Mg^2+^ would decrease the likelihood that the two stems will interact to promote catalysis. These findings provide definitive evidence that both the junction conformer and helix orientation undergo conformational change in the presence of Mg^2+^.

In contrast with evidence for an equilibrium distribution between conformers *in vitro*, recent results from biochemical structure probing in human cells (Anokhina et al. 2013) and by cryo-EM studies of human (Bertram et al. 2017b) and yeast (Yan et al. 2015; Rauhut et al. 2016; Plaschka et al. 2017; Wan et al. 2019) spliceosomes illustrate that the more open three-helix junction is generally observed in the context of intact spliceosomes.

While these studies focused on behavior of junction alone, parallel experiments on complexes in which the ACAGAGA loop was restored, with or without pairing of the U2 snRNA side of the ACAGAGA loop with a short fragment to form a branch site helix, found no detectible difference in the distance between the termini of Helix III and the ISL, even in the presence of high concentrations of Mg^2+^.

These data suggest that although the junction acts as a Mg^2+^-sensitive pivot in positioning the stems, it alone does not drive the proximity of the stems needed for (triplex formation) interaction. The four-helix fold is very far from achieving the anticipated active conformation involving triple helix formation between sequence elements of the ISL, ACAGAGA loop, and AGC triad, with a partial approach of the critical stems upon addition of Mg^2+^; however, the observation of only the three-helix conformer *in situ* may support the hypothesis that it exists to protect premature formation of the active site, or simply that it is simply a lower energy conformer *in vitro* (Sashital et al. 2004). In contrast, the three-helix conformer of the junction observed *in situ* facilitates positioning of stems containing catalytically essential components, but the presence of Mg^2+^ actually drives the stems apart.

Inclusion of the region of the ACAGAGA sequence that participates in the long-range interaction, in the protein-free system, is insufficient to promote folding. This observation suggests that although all elements required for catalytic activity are present in the RNA, the inability to form or stabilize the active site by the U2-U6 snRNA complex alone is mirrored by the exceptionally slow (and low-yield) rate of catalysis by the RNA alone, and even then only a modified reaction by sequences mutated to stabilize interaction (Valadkhan et al. 2009).

These long-range interactions are structurally equivalent to their counterparts in the Group II intron, in which a triple helix is defined by interaction of the Domain 5 bulge and J2/3 with the major groove edge of the catalytic AGC triad (Toor et al. 2008). Crystal structures of a Group IIC self-splicing intron identified the importance of an elaborate network of long-range interactions to stabilize the catalytic DV and other components (Toor et al. 2010); this situation is not paralleled in the U2-U6 snRNA complex in spliceosomes, where regions of snRNAs not directly involved in forming the catalytic site are dispersed in a protein-rich environment (Zhang et al. 2019 and references therein).

The role of RNA-RNA tertiary interactions involving multiple domains to stabilize the catalytic center in the Group II intron is in stark contrast with the reliance on protein-RNA interactions to stabilize the catalytic core of the spliceosome, and points to a vital difference between the two splicing systems.

Protein components clearly fulfill an essential role in facilitating the fold of the U2-U6 snRNA complex into a catalytically active fold, which starts to form in the B^act^ stage, although images (Yan et al. 2016) imply that the catalytic Mg^2+^ ions are not in place until the B* stage. The NTC-related protein RBM22 in the human spliceosome (Cwc2 in *S. cerevisiae*) is implicated in the final folding activity (Rasche et al. 2012; Schmitzová et al. 2012; McGrail et al. 2009). The essential role of Cwc2 in yeast spliceosomes was demonstrated by effects of its deletion from yeast spliceosomes, *i.e.* complete inhibition of the first step of splicing; subsequent supplementation of exogenous Cwc2 rescued the catalytic activity. Rasche *et al*. (2012) have demonstrated these NTC-related proteins make contact with U6 snRNA in the 5′ terminus and the upper part of the ISL – although it was shown that residues of the protein residing in its RRM, finger, and region connecting the Torus and RRM domains may be involved in this binding, it is not clear precisely which region of the protein recognizes which region of the RNA. In other work (Ciavarella, Perea, Greenbaum, unpublished), we have shown that these regions only account for a fraction of binding affinity, suggesting that additional contacts not detectable by cross-linking are important for interaction, and perhaps for their role in RNA folding.

Although the folded state of the U2-U6 snRNA complex starts to form in the B^act^ stage, we anticipate that binding of the RBM22 (or Cwc2), alone or in association with NTC proteins, in addition to anchoring distant regions of the RNA complex, acts as a scaffold for the central junction and precise positioning of the catalytic Mg^2+^. We also speculate that RBM22 preferentially binds the RNA complex in its open three-helix form (already somewhat favored by presence of Mg^2+^ under cellular conditions); binding to the fraction in the three-helix conformer then shifts equilibrium values of the two junction conformers to favor the open three-helix form observed in active spliceosomes, and that this protein-RNA interaction stabilizes formation of the catalytically active form and positioning of divalent metal ions to interact with scissile phosphates at the 5′ splice site.

## MATERIALS AND METHODS

### Design and Synthesis of samples

To determine three-dimensional conformations of stems surrounding the central junction of the protein-free U2-U6 snRNA complex, we first designed, prepared, and paired RNA oligomers labeled with fluorescent dyes at designated sites.

To create a pairing representing the human U2-U6 snRNA complex with the native (WT) sequence in the region of the central junction that would be amenable to measurement of distances between helix termini by FRET, we designed two strands: 1) a 32 nucleotide truncated U6 strand started from the 5′ terminal end of U6 snRNA (*i.e.* starting at G_33_, 8 nucleotide upstream of the ACAGAGA sequence), and terminated just before the hairpin pentaloop of the ISL, synthesized by IDT; and 2) a U6-U2 “chimeric” oligomer starting immediately 3′ of the ISL hairpin loop and connecting with the 5′ stem of U2 snRNA at the end of Helix II *via* a GCAA tetraloop to the U2 snRNA strand through U_46_ of the native sequence. The native sequence at the 5′ terminus of the human U6 snRNA oligomer includes nine nucleotides capable of forming Watson-Crick base pairs with complementary nucleotides at the 3′ end of the U2 snRNA oligomer. Although we observe these base pairs *in vitro* (Zhao et al. 2013), this pairing has been shown not to occur *in vivo* (Anokhina et al. 2013) but since the duplex formation observed in the absence of other spliceosomal components is useful for measurement of FRET-based distances between termini of stems, and thus of angles, between Helix I and the U6 ISL, we have maintained such a duplex in each of our constructs.

To create a construct that focused solely on the native junction without the added flexibility in the Helix I/III stem from the large open ACAGAGA region, we also modified the sequence of U2 snRNA opposing the ACAGAGAA loop of U6 snRNA to form a complementary Helix I/III stem (WTΔL; Figure 1A). All nucleotides within the region forming the central junction are present in the wild type sequences; modifications include truncation of sequence regions beyond paired stems and, in some cases, removal of the ISL pentaloop, a change in the 3′ terminal nucleotide of the truncated U6 strand (U_64_) and the 5′ nucleotide of the chimeric U6/U2 strand (A_70_) to create a C-G pair (maintaining base pairing adjacent to the deleted hairpin loop of U6 ISL) and facilitate transcription of the chimeric strand. The chimeric strand was prepared by *in vitro* transcription techniques using double-stranded DNA templates (IDT) and T7 polymerase overexpressed and purified in the laboratory, followed by gel purification and electroelution of the final product.

We next used a construct, based on WTΔL, that we had previously designed (Zhao et al. 2013) to limit junction conformation to the four-helix conformer by mutating several U2 Stem I nucleotides: mutating G12-C21 to C12-G21, as well as U_17_U_18_ to C_17_G_18_ (Construct 4HJΔL1). These modifications disrupt base pairing of the AGA triad in U6 snRNA to U2 snRNA (disfavoring the three-helix fold) and stabilize Stem I (to favor the four-helix fold), respectively. All other modifications described in WTΔL, including the closing of the ACAGAGA loop, were maintained (Figure 1B).

To measure the distance between the U6 ISL and Helix II while maintaining the same junction features as in 4HJΔL1, we made changes to the design of 4HJΔL1 to enable attachment of linkers and dyes to the 5′ terminus of U2 snRNA (4HJΔL2; Figure 1C). In design of this construct, the connecting tetraloop GCAA was “moved” from the end of Helix II to the end of Helix I/III, resulting in a chimeric strand starting with the 5′ terminus of U2 snRNA, and connecting to the 5′ end of U6 snRNA by the tetraloop. We also removed five base pairs from Helix III (this helix was not dye-labeled in this experiment) to compensate for addition of the loop and added two base pairs to U6 ISL to increase dye-labeling yield. The 5′ 31 nucleotide fragment representing the 5′ side of the ISL and U6 strand of Helix II as synthesized by Dharmacon (synthesized with a 5′ phosphate to facilitate linker attachment); the longer chimeric strand representing U2 snRNA and the 5′ segment of U6 snRNA was generated by *in vitro* transcription as described above.

To measure the distance between 5′ of U2 snRNA and 5′ of U6 snRNA, we designed a construct favoring four-helix junction fold used in Construct 4HJΔL3 (Figure 1D). The mismatch (U6)U_90_-(U2)U_10_ in Helix II was modified to form a Watson-Crick pair by changing (U2)U_10_ to (U2)A_10_; Helix II and Helix III have the same number of base pairs as 4HJΔL1.

To measure the distance between 5′ of U6 snRNA and internal U_16_ of Stem I, we designed construct 4HJΔL4 (Figure 1E). The U6 strand, which represents the U6 snRNA sequences between Helix III and Helix II, including the ISL pentaloop, was transcribed and purified as above. The U2 strand containing a linker-labeled internal U_16_ was synthesized chemically (IDT); U_16_ was incorporated as the amino-modified phosphoramidite iUniAmM. The six-carbon linker is attached covalently through the C5 of uridine, providing a free primary amine that attaches to a fluorophore by the same reaction as attachment at the terminus of an oligomer.

Finally, to assess the impact of the ACAGAGA loop on angles between Helix I/III and the ISL as a result of flexibility or long-range interactions with elements in the ISL, we reintroduced the native U2 snRNA sequence opposing the U6 ACAGAGA sequence in two constructs, WTΔL and 4HJΔL1 to create WT and 4HJ, respectively (Figure 1F and 1G).

All transcribed RNAs were purified by ethanol precipitation and electrophoresis on a 20% polyacrylamide gel. Desired RNA bands were eluted with an Elutrap device at 4 ℃, 100 V overnight. The concentration of RNA was determined by absorbance at 260 nm.

### Pairing of strands

The base pairing of all RNA constructs was performed mixing and annealing 40 pmol of each RNA strand in a buffer of Tris (30 mM)-HEPES (60 mM), pH 7.6, 30 mM NaCl, 1 mM EDTA, by heating at 85℃ for 3 minutes and cooling at room temperature for 10 minutes prior loading on the gel. As controls, individual strands were subjected to the same process. Samples were then applied to nondenaturing 15% PAGE at 4 ℃, 120 V, for 4 hours in a buffer of Tris-HEPES, pH 7.6. Gels were stained with ethidium bromide and visualized under UV light at 302 nm (and/or visualized by fluorescence of attached dyes). Complete pairing was confirmed by appearance of a single band with slower electrophoretic mobility (Figure 2) than either of the individual strands.

### Linker and dye labeling

The donors (fluorescein NHS ester, AF488 NHS ester) and the acceptors (Cy3 NHS ester, AF555 NHS ester) were attached covalently to linker-modified 5′ phosphate termini or a linker-modified internal uridine residue (*e.g.* construct 4HJΔL4) via a two-carbon (in the transcribed RNAs) or six-carbon (in the chemically synthesized RNAs) primary diamine linker. The two-carbon primary diamine linker was added to 5′ phosphate termini using protocol Tech tip #30 provided by Thermo Scientific (ThermoFisher.com). An excess of fluorophore (250 µg dissolved in 14 µl DMSO) was added to 100 µg of amine-modified RNA in 0.1M sodium tetraborate buffer, pH 8.5, to a total volume of 100 µl in a darkened room. The reaction mixtures were mixed occasionally during first two hours and then left to incubate overnight at room temperature. The dye labeled RNAs were purified by ethanol precipitation followed by 20% polyacrylamide gel electrophoresis and eluted by a “crush and soak” method. The dye-labeling yield was determined by absorbance of RNA and the fluorophore at 260 nm and the excitation wavelength of each dye, respectively. The base pairing of labeled RNA strands was tested as described above to confirm that labeling did not diminish pairing efficiency.

### FRET measurements

In these studies, we used FRET measurements to determine distances <100 Å from a donor D to an acceptor A attached to two sites within a paired complex. The rate of energy transferred from D to A is measured by the decrease of Donor emission or by decrease in lifetime of the D at the wavelength where only D emits when A presented. Samples for FRET experiments were made using the same solution conditions used for pairing experiments, with the one difference that [RNA] in FRET samples was 150 nM.

Steady state measurements were performed on Fluorolog-3 from Horiba Scientific. We excited fluorescein at 495 nm (5 nm bandwidth) and measured Cy3 emission intensity at 518 nm. RNA strands labeled with D or A only were used as control samples. For time-resolved measurements (also performed on the Horiba instrument), fluorescence decay curves of D in RNA solutions (150 nM) were recorded on a timescale of 20 ns to a total of 10000 counts in the peak. To measure the lifetime of D only, we used control samples in which the dye labeled RNA was paired with the unlabeled RNA (same RNA with the acceptor used for FRET measurements). In FRET samples, D labeled RNA and A labeled RNA were paired with each other. Instrument response function was measured by Ludox scattering solution in water (internal calibration by instrument). For experiments testing the effect of titration of Mg^2+^, each aliquot of Mg^2+^ was assayed on a separate sample.

### Donor decay lifetimes in the absence and presence of acceptor

All measurements were performed at room temperature in Tri-HEPES buffer, pH 7.6. The fluorescence lifetime of the D in the presence or absence of the acceptor should be measured at wavelength where no emission of the acceptor observed. We excited samples at 495 nm and recorded the signal at 518 nm with fluorescein; 490 nm and 525 nm with AF488 dye. The decay curves were fitted with two-exponential equation giving *α_i_* (fractional amplitude associated with each lifetime) and *τ_i_* (lifetime components of D)

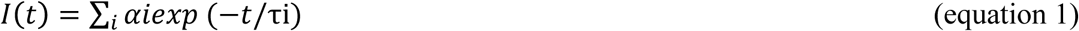

where ∑*αi* is normalized to unity. The results were used to calculate average lifetime of two decay time components, which was used to calculate FRET efficiencies. The average lifetime of D was calculated by 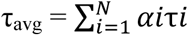. For curves best fit by a biexponential decay, the fitted curve is deconvoluted into two individual lifetimes, τ_1_ and τ_2_, each with its fractional contribution to the total: τ_1_+σ_1_ with relative amplitude of α_1_ and τ_2_+σ_2_ with relative amplitude of α_2_ in which σ*_i_* represents the standard deviation for lifetime *i*. Calculation of the average lifetime of a Donor was obtained by

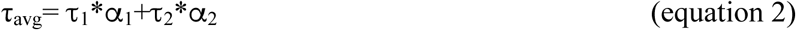

To calculate the standard deviation of the average lifetime, we calculated error propagation by:

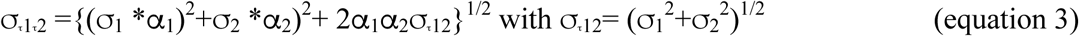

The fitting was judged by reduced chi-squared value (reduced χ^2^), which is expected to be near unity for a good fitting. The FRET efficiency was determined by decrease in lifetime of D in the presence of A (E = 1 – τ_DA_/τ_D_) and results are given in Tables 1.

**Table 1:** Distances between dyes labeled at termini of U2-U6 snRNA constructs used in this study in solution as described in Materials and Methods, without and with 40 mM added Mg^2+^. Constructs WTΔL and WT have the wild type junction sequence that includes a mixture of three- and four-helix conformation. Construct 4HJΔL1, 4HJΔL2, 4HJΔL3, 4HJΔL4 and 4HJ form only the four-helix fold. Distances were calculated using Equation 4, for angles defined by dye placement for each construct (Figure 1), with R_0_ of 56 Å for the fluorescein-Cy3 pair, and R_0_ of 70 Å for the AFF488-AFF555 pair.

| Construct | 0 $Mg^{2+}$<br>Distance Å / (Angle°) | 40 mM $Mg^{2+}$<br>Distance Å / (Angle°) | $\Delta$ Distance (Å) |
| --- | --- | --- | --- |
| WTΔL | 80.0±2.2 Å<br>(84.4±3.6°) | 83.5±1.5 Å<br>(not calculated) | - 3.5 |
| 4HJΔL1 | 90.8±0.6 Å<br>(105.2±1.2°) | 78.3±0.8 Å<br>(81.3±1.4°) | -12.5 |
| 4HJΔL2 | 73.0±1.0 Å<br>(110.1±2.4°) | 67.9±1.0 Å<br>(99.1±2.0°) | -5.1 |
| 4HJΔL3 | 97±1 Å<br>(116.7±2°) | No measurable change | No measurable change |
| 4HJΔL4 | 85.2±0.8 Å<br>(139.0±4.0°) | 87.3±0.8 Å<br>(150.5±5.6°) | +2.1 |
| WT | 84.1±1.4 Å | 75.3±2.6 Å<br>(N/A; mix of angles) | -8.8 |
| 4HJ | 84.0±1.2 Å | 71.3±1.2 Å<sup>†</sup> | -12.7 |
| Calculated 3HJ* | 45.3 Å<br>(19°) | 67.7 Å<br>(63.1°) | +22.4 |

\*Distance and angle between dyes labeled on Helix I/III and U6 ISL in three-helix junction model were calculated from deconvolution for the fraction of four-helix fold (construct 4HJΔL1, which represented ~91.4% and 83.0% of the total in 0 mM Mg<sup>2+</sup> and 40 mM Mg<sup>2+</sup>, respectively) from that of the construct WTΔL. The number in parentheses is the estimated angle between stems. The angle in Construct WTΔL with 40 mM Mg<sup>2+</sup> was not included in the table as it was a meaningless number; a weighted sum of two angles in two conformers.
<sup>†</sup>Angle not reported because change in distance is presumably the result of kinking of the ACAGAGA loop.

### Calculation of Donor-Acceptor Distance

The distance between D and A, *R*, is:

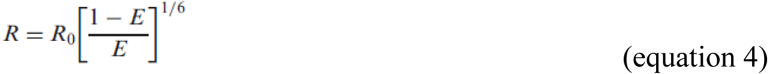

where R_0_ is the Förster distance between D and A at which E (FRET efficiency) is 50%; R_0=_ 56 Å (Norman et al. 2000) for the fluorescein-Cy3 pair and 70 Å for AF488-AF555 pair.

### Calculation of angles between stems

Distances between two termini were translated into estimated angles between stems X and Y. Calculation of stem length assumed A-type helical stacking parameters and included the approximation that the Cy3 dye + linker stacked onto RNA helices, contributing ∼2.6 Å, equivalent to an additional base pair, to the length of a helix. Fluorescein is mobile in solution due to its positive charge (Norman et al. 2000). In order to calculate the angle between two helices, we estimated the overall contribution of fluorescein to the length of RNA helices by performing trFRET on dye-labeled B-type DNA duplexes of 8 and 18 base pairs. Resulting data indicated that fluorescein with a two-carbon primary diamine linker contributes ∼8.5 Å to the length of a duplex.

## ACKNOWLEDGEMENTS

The authors thank Prof. Michael Drain for use of the Horiba Jobin-Yvon fluorometer, Prof. David Lilley for his helpful advice testing dye mobility, Wendy Li and Kenel Zhao for assistance with sample preparation, and Dr. William Perea for a careful reading of the manuscript. NLG acknowledges financial support from the City University of New York PSC-CUNY award program (TRADB-48-370) and from NSF (#MCB-0929394).

